# Comparative genomic analysis of three salmonid species identifies functional candidate genes involved in resistance to the intracellular bacteria *Piscirickettsia salmonis*

**DOI:** 10.1101/589200

**Authors:** José M. Yáñez, Grazyella M. Yoshida, Ángel Parra, Katharina Correa, Agustín Barría, Liane N. Bassini, Kris A. Christensen, Maria E. López, Roberto Carvalheiro, Jean P. Lhorente, Rodrigo Pulgar

## Abstract

*Piscirickettsia salmonis* is the etiological agent of Salmon Rickettsial Syndrome (SRS), and is responsible for considerable economic losses in salmon aquaculture. The bacteria affect coho salmon (CS) (*Oncorhynchus kisutch*), Atlantic salmon (AS) (*Salmo salar*) and rainbow trout (RT) (*Oncorhynchus mykiss*) in several countries, including: Norway, Canada, Scotland, Ireland and Chile. We used Bayesian genome-wide association (GWAS) analyses to investigate the genetic architecture of resistance to *P. salmonis* in farmed populations of these species. Resistance to SRS was defined as the number of days to death (DD) and as binary survival (BS). A total of 828 CS, 2,130 RT and 2,601 AS individuals were phenotyped and then genotyped using ddRAD sequencing, 57K SNP Affymetrix® Axiom® and 50K Affymetrix® Axiom® SNP panels, respectively. Both trait of SRS resistance in CS and RT, appeared to be under oligogenic control. In AS there was evidence of polygenic control of SRS resistance. To identify candidate genes associated with resistance, we applied a comparative genomics approach in which we systematically explored the complete set of genes adjacent to SNPs which explained more than 1% of the genetic variance of resistance in each salmonid species (533 genes in total). Thus, genes were classified based on the following criteria: i) shared function of their protein domains among species, ii) shared orthology among species, iii) proximity to the SNP explaining the highest proportion of the genetic variance and, iv) presence in more than one genomic region explaining more than 1% of the genetic variance within species. Our results allowed us to identify 120 candidate genes belonging to at least one of the four criteria described above. Of these, 21 of them were part of at least two of the criteria defined above and are suggested to be strong functional candidates influencing *P. salmonis* resistance. These genes are related to diverse biological processes, such as: kinase activity, GTP hydrolysis, helicase activity, lipid metabolism, cytoskeletal dynamics, inflammation and innate immune response, which seem essential in the host response against *P. salmonis* infection. These results provide fundamental knowledge on the potential functional genes underpinning resistance against *P. salmonis* in three salmonid species.

## Introduction

Infectious diseases are responsible for large economic losses in salmon farming. *Pisciricketssia salmonis*, the causal agent of Salmon Rickettsial Syndrome (SRS), affects several salmon species and is considered one of the major pathogens affecting the salmon farming industry (Rozas and Enríquez, 2014). *P. salmonis* was identified in 1989 from farmed coho salmon (*Oncorhynchus kisutch*) sampled in Chile (Cvitanich et al., 1991). Since then, *P. salmonis* has been confirmed as the causative agent for clinical and chronic SRS in coho salmon, Atlantic salmon (*Salmo salar*) and rainbow trout (*Onchorhyncus mykiss*) in several countries, including: Norway, Canada, Scotland, Ireland and Chile (Fryer and Hedrick, 2003). Current control protocols and treatments are based on antibiotics and vaccines. The effectiveness of both strategies in field conditions is not optimal (Rozas and Enríquez, 2014). From the total mortalities ascribed to infectious diseases in Chile, SRS is responsible for 18.3%, 92.6% and 67.9% in coho salmon, rainbow trout and Atlantic salmon, respectively (Sernapesca, 2018). These mortality rates, together with other factors such as antibiotic treatments and vaccinations, have generated economic losses up to USD $ 450 million per year (Camusetti et al., 2015).

A feasible and sustainable alternative to prevent disease outbreaks is genetic selection for disease resistance (Bishop and Woolliams, 2014). The estimated levels heritability for resistance to *P. salmonis* in coho salmon, Atlantic salmon and rainbow trout, range from 0.11 to 0.41 (Bangera et al., 2017; Barría et al., 2018a; Bassini et al., submitted; Correa et al., 2015; Yáñez et al., 2016; Yoshida et al., 2018a); demonstrating the feasibility of improving *P. salmonis* resistance through artificial selection in farmed salmon species.

Currently, the advancement of molecular technologies has allowed the generation of dense marker panels for salmonid species (Houston et al., 2014; Macqueen et al., 2017; Palti et al., 2015; Yañez et al., 2016). The use of genotypes from dense panels of SNP markers, together with phenotypes for the traits of interest, assessed in a large number of individuals could provide opportunities to discover the genetic architecture of complex traits. When genetic markers are linked to a major effect quantitative trait loci (QTL), marker assisted selection (MAS), could then be implemented into breeding programs. For instance, a QTL explaining ∼80% of the genetic variance for resistance to Infectious Pancreatic Necrosis Virus (IPNV), has been identified in Scottish and Norwegian Atlantic salmon farmed populations (Houston et al., 2008; Moen et al., 2009). To date, the number of IPN outbreaks has been significantly reduced in Norwegian Atlantic salmon populations because of MAS for IPNV resistance (Hjeltnes, 2018). Interestingly, Moen et al., (2015) mapped the QTL to a region containing an epithelial cadherin (*cdh1*) gene encoding a protein that binds to IPNV, indicating that the protein is part of the machinery used by the virus for host internalization.

*P. salmonis* resistance has been suggested to be polygenic, with many loci explaining a small amount of the total genetic variance (Barría et al., 2018a; Correa et al., 2015), suggesting that the implementation of genomic selection (GS) is the most appropriate strategy to accelerate the genetic progress for this trait. Methods which can model all available SNPs simultaneously, including Bayesian regression methods (Fernando & Garrick 2013), appear to be better for estimating marker effects than conventional methods of modeling each SNP individually, and therefore are becoming increasingly more popular for GWAS (Goddard et al. 2009).

Regarding that *P. salmonis* affects farmed populations of three phylogenetically related salmonid species, including coho salmon, Atlantic salmon and rainbow trout, generating mortalities in a similar manner and that genetic variation for *P. salmonis* resistance has been already reported, we believe that exploring the genetic architecture of this trait simultaneously in the three species can provide further insights into the biology of the differential response against this intracellular bacteria among individuals. Thus, a comparative genomics approach aiming at evaluating and comparing genomic regions involved in *P. salmonis* resistance in coho salmon, Atlantic salmon and rainbow trout would help in narrowing down the list of potential candidate genes associated with the trait for further functional validation in salmonid species.

The aims of this study were i) to dissect the genetic architecture of resistance to *P. salmonis* in coho salmon, Atlantic salmon and rainbow trout using SNP and phenotype data modeled together using Bayesian GWAS approach, ii) to identify genomic regions involved in *P. salmonis* resistance among the three salmonid species and iii) to identify candidate genes associated with *P. salmonis* resistance through a comparative genomics analysis.

## Material and methods

### Challenge tests

A total of 2,606, 2,601 and 2,416 fish belonging to 107, 118 and 105 full-sib families from coho salmon (CS), Atlantic salmon (AS) and rainbow trout (RT), respectively, were independently challenged with an isolate of *P. salmonis* (strain LF-89) (Mandakovic et al., 2016) as described in Barría et al. (2018a), Bassini et al. (submitted) and Yáñez et al. (2013, 2014, 2016). Prior to the beginning of each experimental challenge, qPCR was performed in a sub-sample of each population to confirm the absence of *Flavobacterium* spp, Infectious Salmon Anemia Virus (ISAV) and IPNV. Subsequently, fish were intraperitoneally (IP) injected with 0.2 ml of a LD_50_ inoculum of *P. salmonis*. Post IP injection, infected fish were equally distributed by family into three different test tanks. Each challenge was maintained until mortalities returned to baseline levels. At the end of the challenges, all surviving fish were anesthetized and euthanized. A sample of caudal fin was taken from each survivor and dead fish from each of the experimental challenges for DNA extraction. Body weight was measured at the beginning of the challenge and at the time of death for each individual. The presence of *P. salmonis* was confirmed in a random sample of dead fish through qRT-PCR and necropsy. Each experimental challenge was performed at Aquainnovo’s Research Station, Xth Region, Chile.

### Genotyping

A total of 828 CS, 2,130 RT and 2,601 AS were genotyped using ddRAD, 57K SNP Affymetrix® Axiom® and 50K Affymetrix® Axiom® SNP panels, respectively. Total DNA was extracted using commercial kits following the manufacturer’s protocols. For CS, we used the Wizard SV Genomic DNA purification System (Promega), while DNeasy Blood & Tissue (Qiagen) was used for RT and AS.

For CS, ten double digest Rad-seq (ddRAD) libraries were prepared following the protocol proposed by Peterson et al., (2012), and sequenced on an illumina Hiseq2500 (150 bp single-end). Raw sequences were analyzed using STACKS v. 1.41 (Catchen et al., 2011, 2013). Rad-tags which passed the *process_radtags* quality control were aligned to the coho salmon reference genome (GCF_002021735.1). Loci were built with *pstacks* setting a minimum depth coverage of three. After catalog construction, rad-tags were matched using *sstacks*, and followed by *populations* using default parameters. Quality control (QC) included the removal of SNPs below the following thresholds: Hardy-Weinberg Equilbrium (HWE) p<1×10−6, Minor Allele Frequency (MAF) < 0.05, and genotyping call rate < 0.80. Individuals with a call rate below 0.70 were removed from the subsequent analysis. For a detailed protocol of library construction and SNP identification see Barría et al. (2018a).

RT individuals were genotyped using the commercial 57K Affymetrix® Axiom® SNP array, developed by the National Center of Cool and Cold Water Aquaculture at the USDA (Palti et al., 2015). SNPs were filtered with the following QC parameters: HWE p < 1×10−6, MAF < 0.05, and SNP call rate < 0.95. Individuals with call rates lower than 0.95 were also removed.

The 50K SNP Affymetrix® Axiom® array used to genotype AS, was developed by Universidad de Chile and Aquainnovo (Correa et al., 2015; Yañez et al., 2016). These markers were selected from a 200K array, as described in detail in Correa et al. (2015). Genotypes were quality-controlled using the following criteria: HWE p < 1×10−6, MAF < 0.05, SNP and samples were discarded when the genotype rate was < 0.95.

### Genome-wide association analysis

Resistance to SRS was defined as both the number of days to death (DD) post experimental challenge and as binary survival (BS; 0 for surviving individuals at the end of the experimental challenge and 1 for deceased fish). The GWAS analyses were performed using the Bayes C method which assumes distributed mixture distribution for marker effects. All model parameters are defined in the following equation:

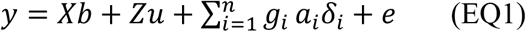

where, y is the vector of phenotypic records (DD or BS); X and Z are the incidence matrix of fixed effects and polygenic effect, respectively; b is the vector of fixed effects (tank and body weight); u is the random vector of polygenic effects of all individuals in the pedigree; *g_i_* is the vector of the genotypes for the *i^th^* SNP for each animal; *a_i_* is the random allele substitution effect of the *i^th^* SNP; *δ_i_* is an indicator variable (0, 1) sampled from a binomial distribution with parameters determined such that π value of 0.99; and *e* is a vector of residual effects.

The prior assumption is that SNP effects have independent and identical mixture distributions, where each SNP has a point mass at zero (with probability π) and a univariate Gaussian distribution (with probability 1-π) with a mean equal to zero and variance equal to 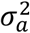, having in turn a scaled inverse *χ*^2^ prior, with *v_a_* = 4 and *v_e_* = 10 degrees of freedom (df) and scale parameter, respectively (Fernando and Garrick, 2013). These hyperparameter values were chosen based on previous studies (Peters et al., 2012; Santana et al., 2016; Wolc et al., 2016; Yoshida et al., 2017, 2018a).

The analyses were performed using the GS3 software (Legarra et al., 2013). A total of 200,000 iterations in Gibbs sampling were used, with a burn-in period of 20,000 cycles and the results were saved every 50 cycles. Convergence was assessed by visual inspection of trace plots of the posterior density of genetic and residual variances.

The proportion of the genetic variance explained by each significant SNP was calculated as:

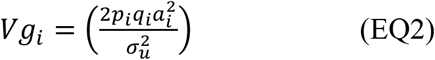

where *p_i_* and *q_i_* are the allele frequencies for the i-th SNP, *a_i_* is the estimated additive effect of the i-th SNP on the phenotype and 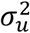 is the estimate of the polygenic variance (Lee et al., 2013).

The association between the SNPs and the phenotypes was assessed using the proportion of the genetic variance explained by each marker. To be inclusive regarding the genomic regions to be compared across the three species, we selected each of the regions explaining at least 1% of the genetic variance for the trait in each species.

The heritability values were calculated as:

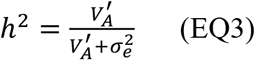

where, 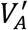 is the total additive genetic variance, estimated as the sum of additive marker 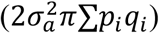 and the polygenic pedigree based 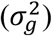 additive genetic variance.

### Comparative genomic analysis

Initially, sequence homology between chromosomes containing regions with SNPs explaining more than 1% of the genetic variance were compared. Synteny among these chromosomes was identified by using Symap (Soderlund et al. 2011). The relationship between the chromosomes from CS, AS and RT and the association between SNPs and resistance to *P. salmonis* (Manhattan plot) was plotted using Circos (Krzywinski 2009).

To identify candidate genes associated with *P. salmonis* resistance, we used a comparative genomic analysis between coho salmon CS, AS and RT. For this, we mapped the location of each SNP that explained 1% or more of the genetic variance for the trait on the reference genome (NCBI_RefSeq) of each species; CS (GCF_002021735.1), AS (GCF_000233375.1, Lien et al., 2016) and RT (GCF_002163495.1, Pearse et al., bioRxiv). Subsequently, we retrieved the sequences of all the genes (and their protein products) adjacent to each SNP within a window of 1 Mb (500 Kb downstream and 500 Kb upstream to the associated SNP). We then used this information to apply the following criteria in order to classify and prioritize functional candidate genes by comparing the genomic regions involved in *P. salmonis* defined as DD and BS within and among the three species:

i. The complete set of genes were identified and classified into homologous superfamilies based on InterPro (Mitchell et al, 2019) protein domain signatures using Blast2GO software version 5.2.5 (Götz et al, 2005) (referred to as: Group A);
ii. Orthologous and paralogous genes among species were identified using the ProteinOrtho tool (Lechner et al., 2011). Multi-directional alignments were performed using the full-length sequences among complete sets of proteins encoded in each of the three species to obtain orthologous groups, with a 35% threshold for identity and similarity (Group B);
iii. The complete set of genes within 1 Mb windows adjacent to SNPs explaining the highest proportion of the genetic variation for each trait (leader SNP) were recovered and classified as high priority genes (Group C); and
iv. The complete set of genes located at the intersection of more than one 1 Mb windows within a species were also identified and considered as high priority genes (Group D).

## Results

### Challenge test and genetic parameters

There was considerable phenotypic variation for *P. salmonis* resistance across fish species (Figure 1). The average cumulative mortality for different families ranged from 5% to 81%, 8.3% to 73.7% and 8% to 100% for CS, AS and RT, respectively. This result suggests that the phenotypic variation for this trait could be related with the genetic background on each species. Estimated heritabilities for *P. salmonis* resistance were significant for the three species, indicating the feasibility to improve the trait by means of artificial selection (Table 1). The genomic heritability values for DD were 0.32 for CS, 0.24 for AS and 0.48 for RT. When resistance was defined as BS, genomic heritability estimates increased to 0.88, 0.32 and 0.64 for CS, AS and RT, respectively, representing moderate to high levels of genetic variation for *P. salmonis* resistance.

**Figure 1.**
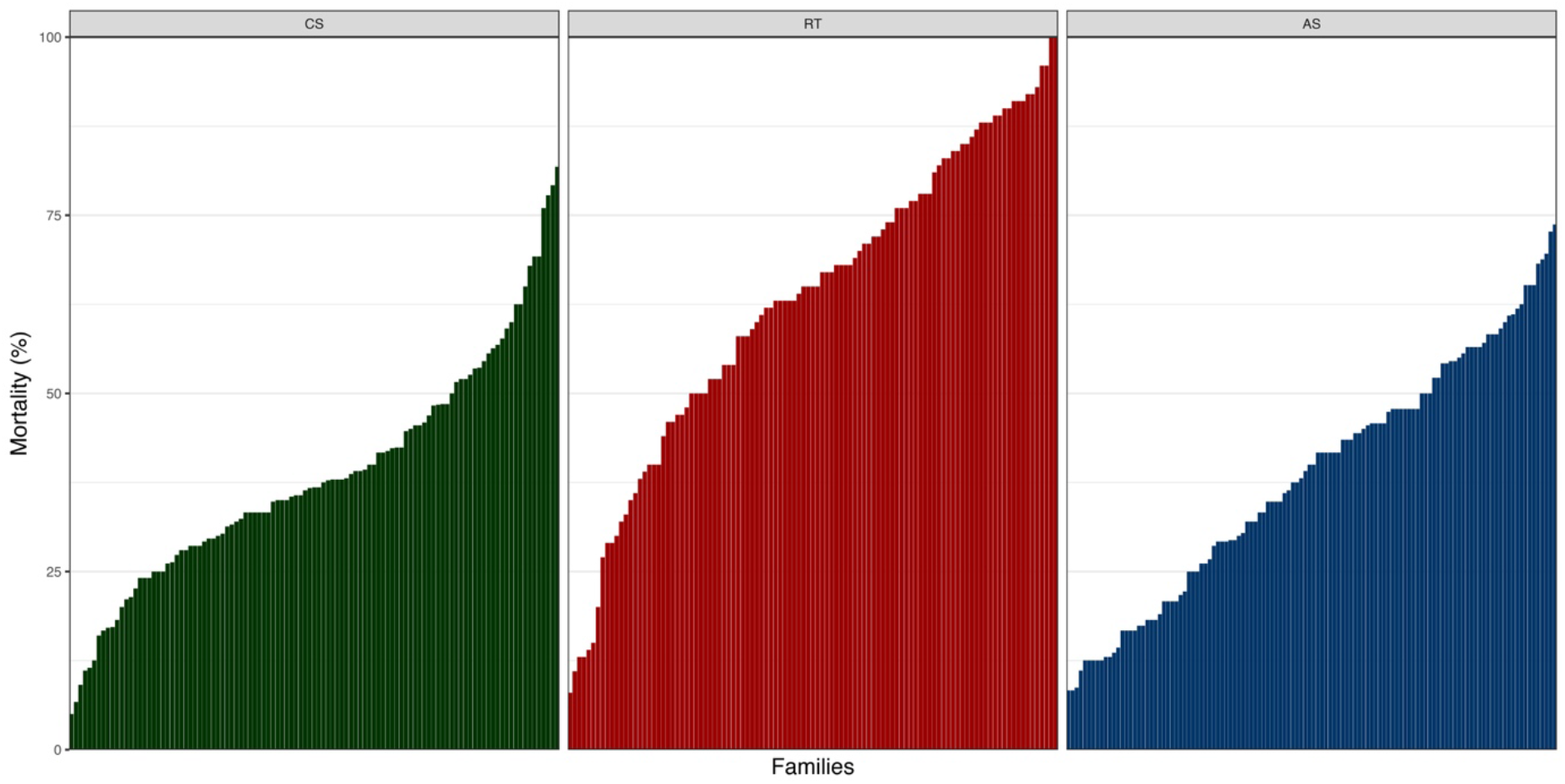
Cumulative mortality by family after *Piscirickettsia salmonis* experimental infection of coho salmon (CS), rainbow trout (RT) and Atlantic salmon (AS). For CS, RT and AS a total of 107, 105 and 118 full-sib families were experimentally challenged.

**Table 1.** Estimates of total additive genetic variance (*V*′*_a_*), residual variance 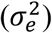, heritability (h^2^) and standard deviation (SD) for resistance against *Piscirickettsia salmonis* in three salmonids species.

| Species | Days to death |  |  |  | Binary survival |  |  |  |
| --- | --- | --- | --- | --- | --- | --- | --- | --- |
| | $V'_a$ | $\sigma_e^2$ | $h^2$ | SD | $V'_a$ | $\sigma_e^2$ | $h^2$ | SD |
| Coho salmon | 28.91 | 60.70 | 0.32 | 0.07 | 7.53 | 1.00 | 0.88 | 0.03 |
| Rainbow trout | 30.42 | 32.71 | 0.48 | 0.04 | 1.87 | 1.00 | 0.64 | 0.05 |
| Atlantic salmon | 16.52 | 53.17 | 0.24 | 0.04 | 0.47 | 1.00 | 0.32 | 0.05 |

### Genome-wide association analysis

A total of 580 CS (9,389 SNPs), 2,383 AS (42,624 SNPs) and 1,929 RT (24,916 SNPs) were retained after QC. For CS and RT we found relatively few SNPs explaining a moderate to high percentage of genetic variance for *P. salmonis* resistance. In contrast, for AS, a large number of SNPs with small effect were found, and the percentage of genetic variance explained by a single marker was not higher than 5% (Figure 2 **and** Supplementary Figure 1). While there were multiple shared syntenic regions with associated SNPs (4 for DD, and 5 for BS) in two species, there were no shared syntenic regions where all three species had common associated SNPs (Figure 2). Figure 3 (**and** Supplementary Figure 2) highlights the different genetic architecture for resistance to *P. salmonis* among the three salmonid species studied. For CS and RT, both traits appear to have oligogenic control with few moderate to large effect loci, and a large-unknown number of loci each having a small effect on the traits. For BS, the top 200 SNPs explained about 80% and 90% of the phenotypic variance in CS and RT, respectively, while in AS they explained slightly more than 30%. For DD, the top ten SNPs explained a 40%, 57% and 17% of the total genetic variance for *P. salmonis* resistance in CS, RT and AS, respectively.

**Figure 2.**
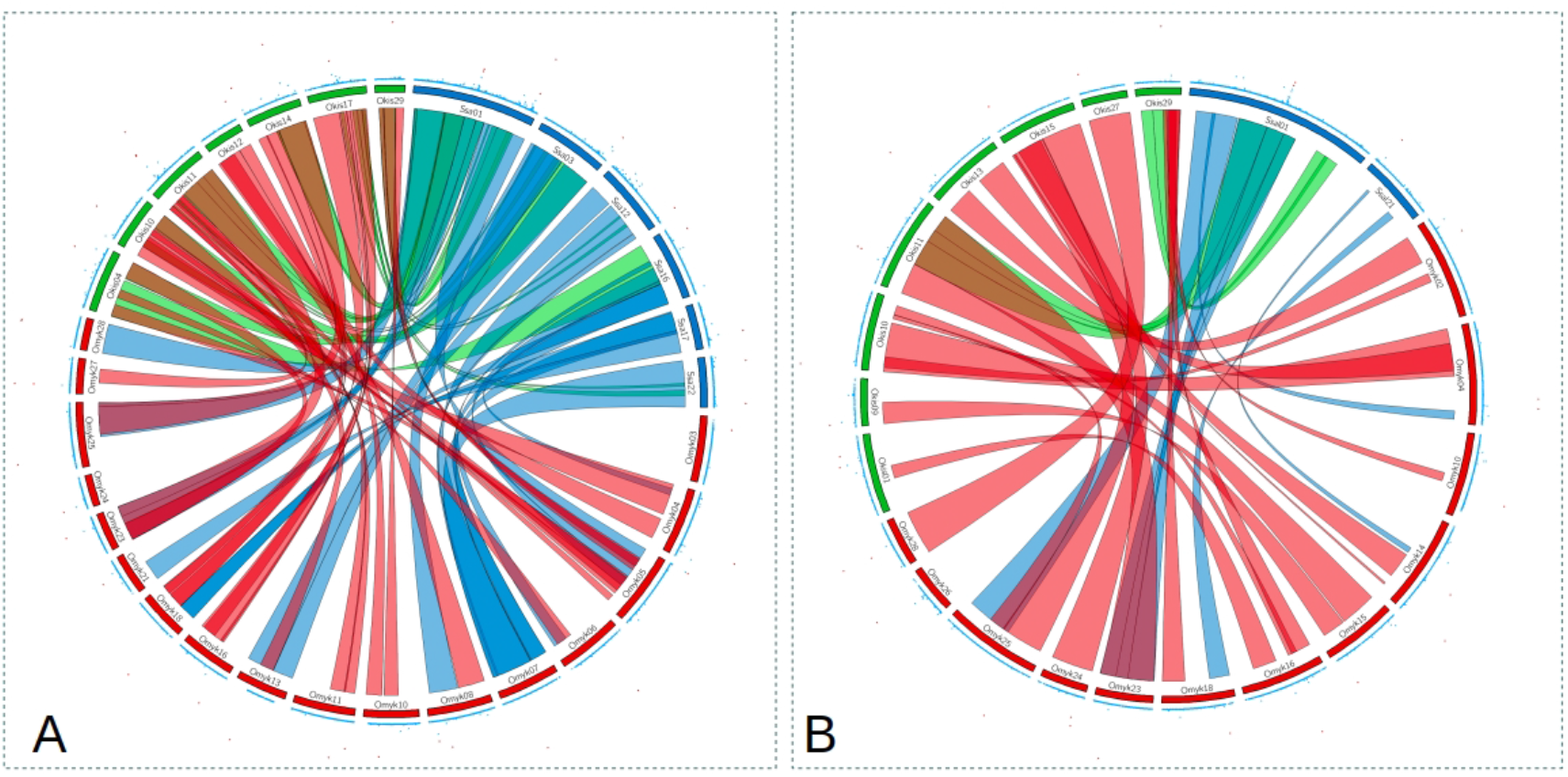
Circos plot for *P. salmonis* resistance as day of death (A) and as binary survival (B). The inner ribbons mark syntenic regions between coho salmon (green) rainbow trout (red) and Atlantic salmon (blue). Manhattan plots are showed on the outer ring, with significant associations plotted in red (values >= 1).

**Figure 3.**
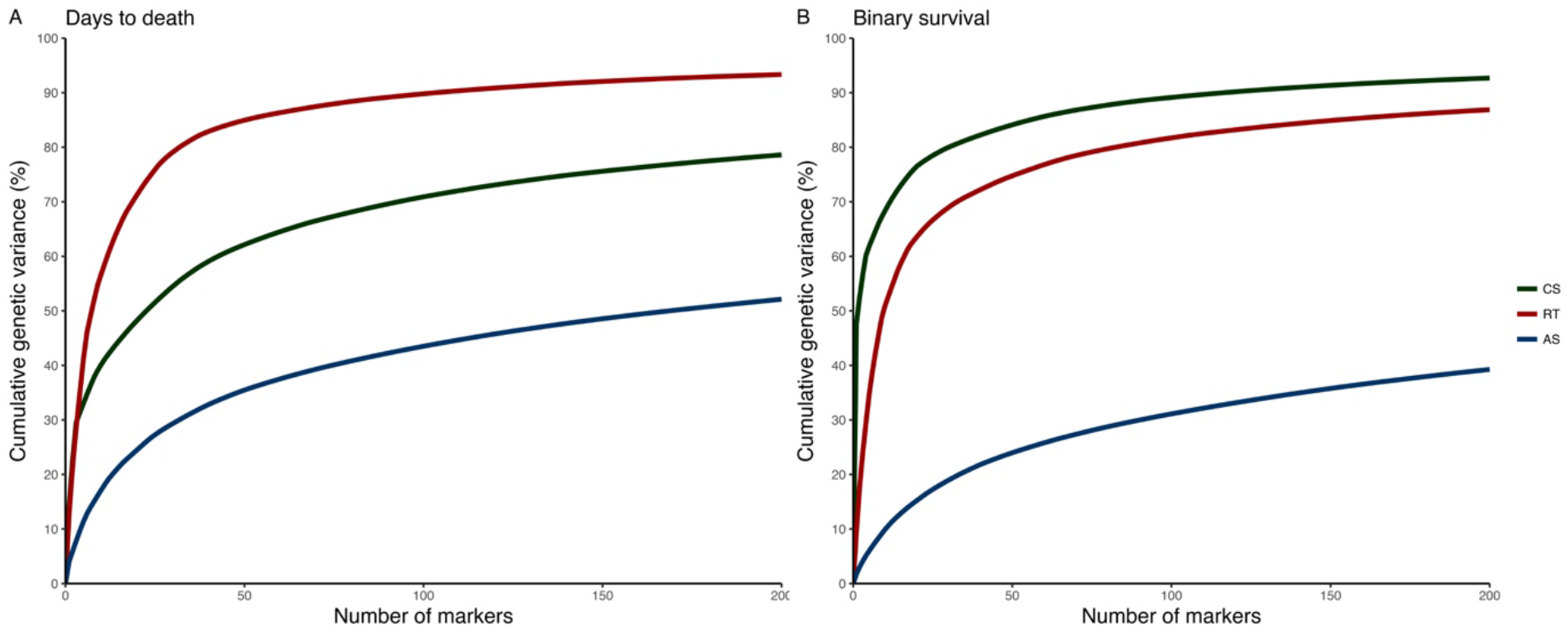
Cumulative percentage of the genetic variance explained (GEV) by the top 200 markers from Bayesian GWAS for resistance to *P. salmonis* measured as days to death (DD) (A) and binary survival (BS) (B) in Coho salmon (CS), Rainbow trout (RT) and Atlantic salmon (AS).

### Comparative genomic analysis

We mapped the location of each SNP that explained 1% or more of the genetic variance for both DD and BS, to the reference genome of CS, AS and RT, and searched for genes within 1Mb windows flanking each SNP. This search allowed us to identify 533 unique genes that encoded 957 proteins. The complete list of genes and proteins can be found in the **Supplementary Table S1: Sheets 1 to 6.**

To prioritize functional candidate genes, we annotated and classified the complete set of encoded proteins in homologous superfamilies for each trait and species, based on InterPro protein domain signatures. We identified 194 and 129 homologous superfamilies for DD and BS, respectively, 103 of which were shared between traits (Supplementary Table S1: homologous superfamilies). The homologous superfamilies and the number of proteins present in at least two salmonids species are shown in Figure 4. Remarkably, around 30% of the proteins from genes present in regions associated with DD belong to five homologous superfamilies (*P-loop containing nucleoside triphosphate hydrolase, immunoglobulin-like fold, zinc finger C2H2 superfamily, zinc finger, RING/FYVE/PHD-type and protein kinase-like domain superfamily*). A total of 30% of proteins from genes present in regions associated with BS belong to only three homologous superfamilies (*P-loop containing nucleoside triphosphate hydrolase, immunoglobulin-like fold and immunoglobulin-like domain superfamily*). Interestingly, the *P-loop containing nucleoside triphosphate hydrolase* superfamily (also known as P-loop_NTPase) contained the largest group of proteins for both traits, and at least one representative protein from each salmonid species belonged to this superfamily. Thirty-one of the proteins identified in this study are part of this superfamily, including some GTPases, kinesin and myosin proteins and ATP-dependent RNA helicases (**Supplementary Table S1, sheet: P-loop NTPases (Group_A)**).

**Figure 4.**
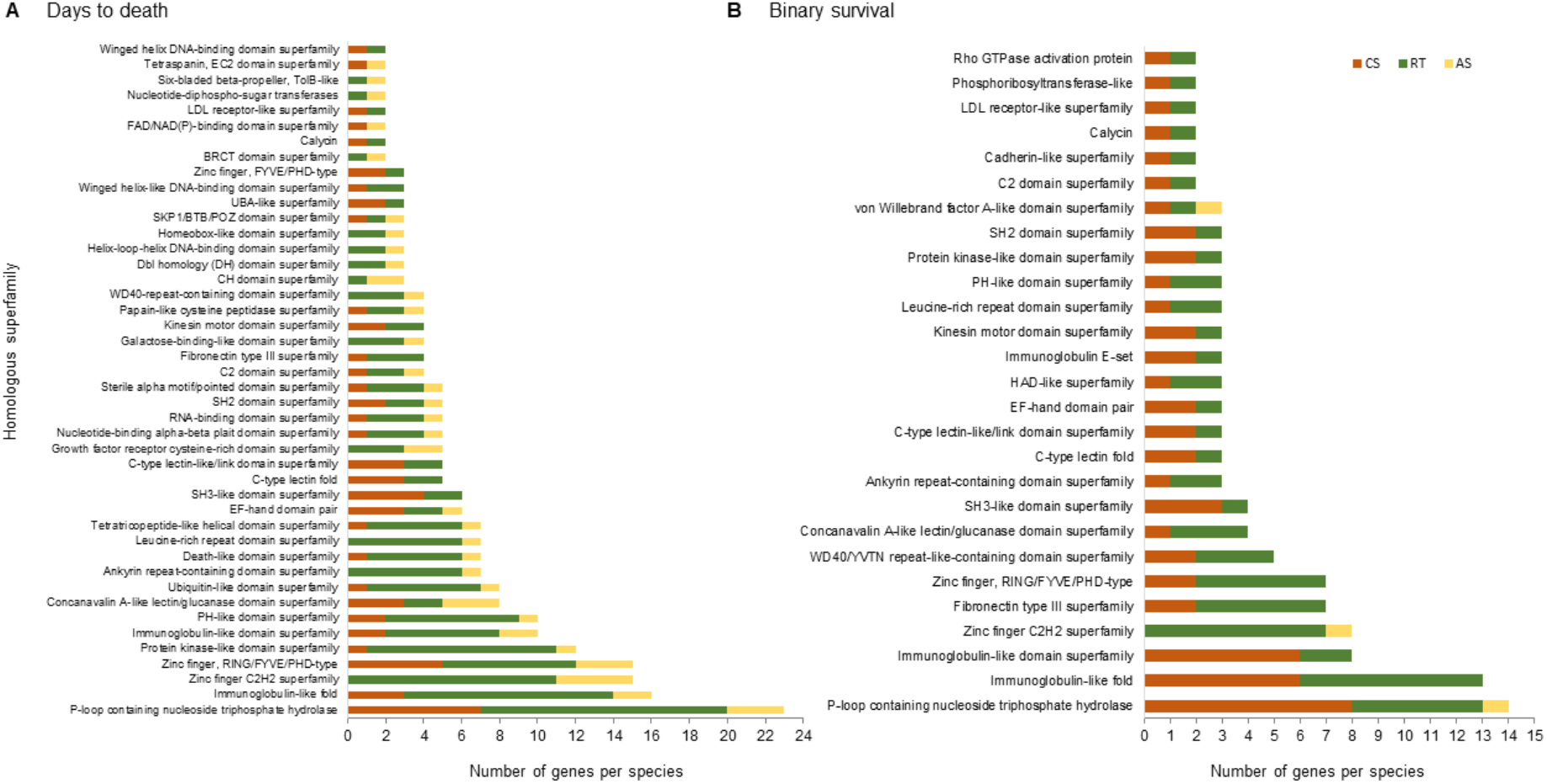
Homologous superfamilies (InterPro) adjacent to the complete set of SNPs that explain over 1% of the genetic variance of resistance to SRS measured as days to death (DD) (A) and binary survival (BS) (B). Bars represent the abundance of genes in each homologous superfamily present in at least two salmonids species. Coho salmon (CS), rainbow trout (RT) and Atlantic salmon (AS).

To complement these analyses, we looked for orthologous proteins through multi-directional alignments using full-length sequences of the complete set of proteins for each species (Group B). Only five groups of orthologous genes were identified in at least two species, highlighting three non-receptor tyrosine-protein kinases (nr-TPK) with representative genes in the three species for DD and in two species for BS. In addition, for DD, two ATP-dependent RNA helicases (DDX) and two Ras-related proteins (RAB) were identified in CS and RT, while two FYVE, RhoGEF/PH domain-containing proteins (FGD) were identified in RT and AS. For BS, two fatty acid-binding proteins (L-FABP) and two ankyrin repeat domain-containing proteins were identified in CS and RT (**Supplementary Table S1, sheet: Orthologous genes (Group_B)**). The proteins nr-TPK, DDX and L-FABP are also encoded by genes adjacent to SNPs that explained the highest proportion for the genetic variance (leader SNP) for both trait definitions (Group C).

Group C contained other genes (n=42) that encoded proteins such as myosin-IIIb (MYO3B), ATP-dependent RNA helicase (TDRD9), kinesin protein (KIF15) and kinesin protein (KIF2C) that are also included into the P-loop_NTPases superfamily, as well as members of the orthologous groups such as fatty acid-binding proteins (FABP). Other genes encoding proteins classically associated to immune response such as tripartite motif-containing protein 35 (TRIM35) and lysozyme C II (LYZ) are also part of this group. A complete list of these genes and proteins is in **Supplementary Table S1, sheet: Adjacent to leader SNP (Group_C)**.

Group D was composed of genes (n=58) located adjacent to more than one SNP simultaneously (within overlapped windows). Among them, we identified GTPase IMAP family member 4 (GIMAP4), GTPase IMAP family member 8 (GIMAP8), NLR family CARD domain-containing protein 3 (NLRC3), ADP-ribosylation factor protein 5B (ARL5B), voltage-dependent L-type calcium channel subunit beta-2 (CACNB2) and heparan sulfate glucosamine 3-O-sulfotransferase 3A1 (HS3ST3A1), all of which are also P-loop NTPases. In addition, we identified histidine triad nucleotide-binding protein 1 (HINT1), that is also adjacent to the leader SNP for DD in AS, and other genes associated with immune response such as collectin-12 (COL12), macrophage mannose receptor 1 (MRC1) and tapasin-related protein (TAPBPL). A complete list of these genes and proteins can be found in **Supplementary Table S1, sheet: Genes overlapped windows (Group_D)**. Additionally, the gene that codes for NACHT, LRR and PYD domains-containing protein 12 (NLRP12) was found in groups A, C and D.

We identified several candidate genes associated with *P. salmonis* resistance (n=120) which were present in at least one of the groups described previously. These genes are associated with the following biological processes: dependent on kinase activity, GTP hydrolysis, helicase activity, lipid metabolism, cytoskeletal dynamics and inflammation. In order to rank the genes, we scored them based on the counting of each of them across following categories: i) species (CS, RT and AS); ii) trait definitions (DD and BS); and iii) groups (A, B, C and D), thus the maximum score for one particular gene was equal to 9. The prioritized functional candidate genes based on the score described above are shown in Table 2 and the complete list of unique candidate genes (n=120) can be found in the **Supplementary Table S1. sheet: Candidate Genes**.

**Table 2.**
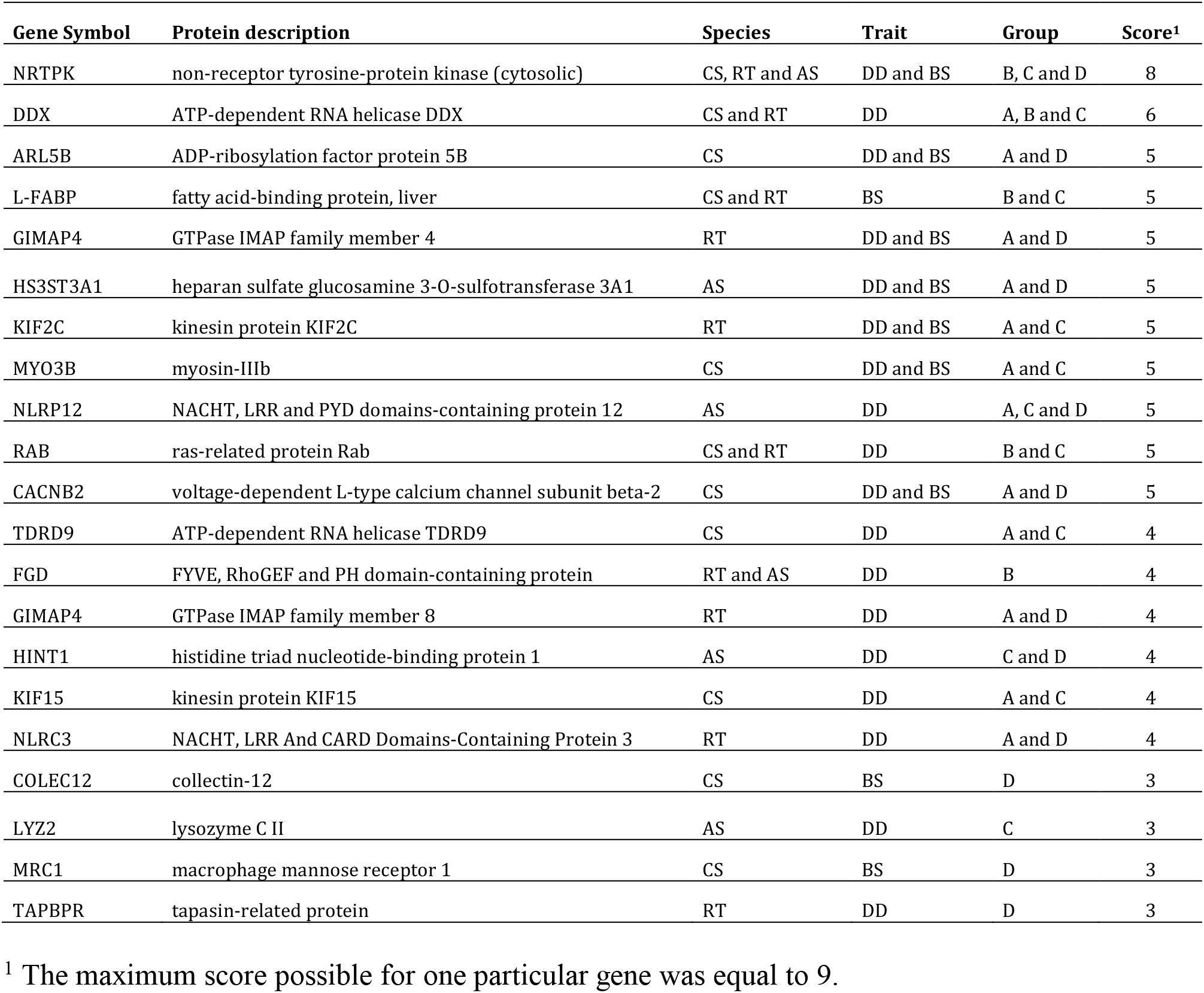
Summary of candidate genes associated with *P. salmonis* resistance for coho salmon (CS), rainbow trout (RT) and Atlantic salmon (AS) ranked by score, which is simply based on the number of appearance of each gene across the following categories: i) species (CS, RT and AS); ii) trait definitions (DD and BS); and iii) groups (A, B, C and D).

## Discussion

The comparative genomic strategy used in this study allowed us to identify groups of homologous superfamilies and orthologous genes common to more than one species of salmonids among genes adjacent to SNPs that explain more than 1% of the genetic variance for *P. salmonis* resistance. To our knowledge, this is the first study which aims at identifying and prioritizing functional candidate genes involved in the differential response against bacterial infection by means of comparing results from genome-wide association mapping across different phylogenetically related salmonid species.

Heritability estimates are in agreement with previous studies aimed to estimate levels of genetic variation for resistance to bacterial diseases in salmonid species. For instance, Vallejo et al., (2016; 2017) presented heritabilities ranging from 0.26 to 0.54 and from 0.31 to 0.48, for resistance to bacterial cold water disease in a farmed rainbow trout population. The levels of genetic variation observed in the current study are consistent or somewhat higher than previous estimates of heritabilities for resistance to *P. salmonis*, depending on the species and the trait definition. For instance, previous heritability values for *P. salmonis* resistance, estimated based on pedigree information reached a maximum of 0.16, 0.41, and 0.44, for CS, AS and RT, respectively (Bassini et al., submitted; Yáñez et. al., 2013, 2014; 2016). When heritability for *P. salmonis* resistance was estimated based on genomic information, the maximum values previously reported were 0.39, and 0.62, for AS and RT, respectively (Bangera et al., 2017; Yoshida et al., 2018a).

Our results show evidence of alleles of medium to large effect involved in resistance to *P. salmonis* in CS and RT. In contrast, for AS our results suggest that if alleles of large effect do exist, they are at such low frequency that they individually explain a small proportion of the variance for resistance to *P. salmonis*. The identification of genomic regions harboring associated SNPs was based on GWAS using the Bayes C approach, which is more suitable for oligogenic traits (Habier et al., 2011). In a few cases, the same SNP was significantly associated with both trait definitions (DD and BS). This could be the result of pleiotropy, closely linked genes (local linkage disequilibrium) or by a strong correlation between both traits. For example, we observed the same SNP associated with DD and BS in CS (58185_41 and 24601_47) and RT (AX-89926208 and AX-89966072) among the top ten SNPs explaining most of the genetic variance for the trait.

Based on the linkage disequilibrium (LD) of the Atlantic salmon population, (measured as r^2^), the number of SNPs used for AS (∼ 43K) should be enough to cover the entire genome (Barría et al., 2018b). There is a lack of studies aimed at evaluating the LD and population structure of the current farmed rainbow trout population. Based on results from a different rainbow trout farmed population, at least 20K SNPs are necessary to cover the whole genome (Vallejo et al. 2018). If LD levels of the present rainbow trout population are similar to those reported by Vallejo et al. (2018), the 23K SNPs used here will most likely cover the whole genome. However, this is not the case for CS. Using a high density SNP array Rondeau et al. (2018, in prep.) and Barría et al. (2018c), suggested that at least 74K SNPs are necessary for whole-genome studies of the current coho salmon population. The small number of SNPs assayed in this study for CS (9,389), most likely affected the identification of markers with a moderate to high effect on resistance to *P. salmonis* in this species.

While the complete set of proteins predicted from reference genomes of CS, RT and AS consisted of 57,592, 58,925 and 97,738, respectively, the proteins neighboring SNPs associated with resistance (range of 1Mb) represent less than 1% of the different proteomes. The characterization of the complete set of proteins among species established that the most prevalent homologous superfamily was the *P-loop containing nucleoside triphosphate hydrolase*. However, since this superfamily contains proteins with at least 21 functions (Shalaeva et al., 2018), it is possible that the high frequency of proteins identified from this group was due to the overall high representation in salmonid genomes. For this reason, we retrieved the sequences of 100 randomly selected proteins from the genomes of CS, RT and AS, and classified them into subfamilies (Supplementary Figure S3). The results indicate that P-loop_NTPase is not the most prevalent in any of the salmonid species, which suggests that this homologous superfamily is actually enriched in the regions analyzed and is not a consequence of their high representation in CS, RT and AS genomes.

When traits are polygenic in nature, the identification of genes underlying them is a challenging task and often depends on previous knowledge of the function of genes adjacent to the associated SNPs (Jiang *et al*., 2014; Bouwman *et al*., 2018; Robledo et al., 2019). Our strategy was based on identifying orthologous proteins between the salmonid species and families of homologous proteins in the complete set of proteins adjacent to all the SNPs that explained more than 1% of the genetic variance, without searching for a specific function. The identification of genes directly associated with the innate immune response, after applying all the classification criteria, such as lysozyme C II, macrophage mannose receptor 1, collectin-12 and tapasin-related protein, suggests that our strategy was successful in finding strong functional candidate genes involved in resistance to *P. salmonis*. Interestingly, around one hundred genes not classically associated with the immune system were also identified; from which seventeen of them were part of at least two of the groups described previously and hence are considered strong candidates for being responsible on trait variation (Table 2).

Previously, lysozymes have primarily been described as having a bacteriolytic activity against gram-positive bacteria; however, the expression of lysozyme C II has been shown to be induced in a resistant rainbow trout line in response to *Flavobacterium psychrophilum* infection (Langevin et al., 2012) and in Atlantic salmon families in response to *Piscirickettsia salmonis* infection (Pulgar et al., 2015), indicating that the transcriptional regulation of this enzyme in salmonids responds to gram-negative bacterial infection. Macrophage mannose receptor 1 and collectin-12 are membrane receptors which display several functions associated with innate immunologic defense, particularly in the recognition of carbohydrate structures of pathogens and as phagocytic receptors of bacteria, yeasts and other pathogenic microorganisms (Harris *et al*., 1992; Ma *et al*., 2015). It has been reported that enhanced infection in human phagocytes with *Francisella tularensis*, a bacterium phylogenetically related to *P. salmonis*, is mediated by the macrophage mannose receptor (Schulert and Allen, 2006), while collectin-12 led to the activation of the alternative pathway of complement via association with properdin, a key positive regulator of the pathway by increment of the half-life of the C3 and C5 convertases (Ma *et al*., 2015). Tapasin-related protein has been described as a second MHC class I-dedicated chaperone, essential to providing specificity for T cell responses against viruses and bacteria (Hermann et al., 2015) and the related protein tapasin has been shown to be induced in monocyte/macrophage in rainbow trout by chum salmon reovirus infection (Sever et al., 2014).

Another set of candidate genes for SRS resistance in the three salmonid species studied are a cluster of cytosolic non-receptor tyrosine-protein kinases (nRTKs). These proteins are a subgroup of the tyrosine kinase family, enzymes that phosphorylate tyrosine residues of proteins, and regulate many cellular functions, such as: cell growth and survival, apoptosis, cell adhesion, cytoskeleton remodeling and differentiation (Neet and Hunter, 1996). Although these genes are not classically related to the response to pathogens, it has been described that the interaction of T- and B-cell antigen receptors with some non-receptor tyrosine protein kinases is critical to the activation of lymphocytes by an antigen (Sefton and Taddie, 1994). Moreover, some cellular signaling pathways are hijacked by intracellular pathogens (bacteria and viruses), thus pathogens can subvert protein phosphorylation to control host immune responses and facilitate invasion and dissemination. It has been described that some bacterial effectors are injected into host cells through their secretion systems where they inhibit the Src kinase (a subfamily of nRTKs). In particular, the effector EspJ, an ADP-ribosyltransferase of the bacteria *Escherichia coli* and *Citrobacter rodentium*, regulates multiple non-receptor tyrosine kinases *in vivo* by ADP-ribosylation, demonstrating that part of its target protein repertoire involves Src kinases such as YES1 and LYN, as well as the adapter SYK (Young et al., 2014; Pollard et al., 2018), all of which were identified in this study in CS, RT and AS. Remarkably, among the candidate genes we also identified the small GTPase ADP-ribosylation factor protein 5B (ARL5B), suggesting that an adequate regulation of the activity of nRTKs by ADP-ribosylation could be critical to combat *P. salmonis* infection.

Other orthologous candidate genes identified in this study encode for proteins RAB1 and RAB18, both members of the GTPase superfamily. The GTPases are a large family of hydrolase enzymes that bind and hydrolyze GTP and play an important role in signal transduction, protein translation, control and cellular differentiation, intracellular transport of vesicles and cytoskeletal reorganization, among other cellular processes (Bourne et al., 1991). Specifically, the RAB GTPases constitute a subfamily of small GTPases known as master regulators of intracellular membrane traffic (Stenmark, 2009). Since *P. salmonis* drives the formation of host membrane-derived organelles, the development of these *P. salmonis*-containing vacuoles (PCVs) are dependent on the bacterium’s ability to usurp the intracellular membrane system of the fish. Interestingly, *Legionella pneumophila*, the closest phylogenetical bacterium to *P. salmonis*, disturbs the intracellular vesicular trafficking of infected human cells by recruiting the RAB1 to the cytosolic face of the Legionella-containing vacuole (LCV) through the activity of its effector protein DRRA (Müller et al., 2010), suggesting that *P. salmonis* could use similar mechanisms for the formation and maintenance of its replicative vacuole. Furthermore, two orthologous of FYVE, RhoGEF and PH domain-containing proteins were identified in RT and AS. These proteins activate CDC42, a GTPase involved in the organization of the actin cytoskeleton and with a role in early contractile events in phagocytes (Ching et al., 2007). Since it has been described that the infective process of *P. salmonis* depends on the exploitation of the actin monomers (Ramírez et al., 2015), the identification in this study of candidate genes that encode for cytoskeletal motor proteins (two kinesins and a myosin), highlights their relevance not only for the reorganization of the cytoskeleton, but also for its motility and involvement in the development of the infection (Hoyt et al., 1997). Remarkably, two other candidate proteins associated with SRS resistance are also members of the GTPase superfamily, the GTPases of the immunity-associated proteins (GIMAPs) 4 and 8. This is a family of proteins abundantly expressed in lymphocytes and whose function is to contribute in the regulation of apoptosis and the maintenance of T-cell numbers in the organism (Yano et al., 2014).

Another group of orthologous genes code for ATP-dependent RNA helicases DDX24 in CS and DDX47 in RT for DD. The ATP-dependent RNA helicase DDX family, also known as DEAD-box helicases, is required for different cellular processes such as transcription, pre-mRNA processing, ribosome biogenesis, nuclear mRNA export, translation initiation, RNA turnover and organelle function. The protein structure is very similar to viral RNA helicases and to DNA helicases, which suggests that the fundamental activities of these enzymes are similar (Rocak and Linder, 2004). Viruses also utilize RNA helicases at various stages of their life cycle. Many viruses carry their own helicases to assist with the synthesis of their genome, but others synthesize their genome within the cell nucleus which tends to exploit cellular helicases and thus do not encode their own. We also identified the ATP-dependent RNA helicase TDRD9, which has not been directly implicated in infection, but was differentially expressed in channel catfish (*Ictalurus punctatus*) in response to *Aeromonas hydrophila* infection (Li et al., 2012). Mechanistic studies of RNA helicases will allow the determination of the precise role of these helicases in the host/pathogen interaction.

The last group of orthologous genes identified code for two liver fatty acid binding proteins (L-FABP) in CS and RT for BS. Liver fatty acid-binding proteins are abundant in hepatocytes and are known to be associated with lipid metabolism. In addition, these proteins are up-regulated in several types of cancer but their role in infection remains unclear (Ku et al., 2016). Nevertheless, it has been recently reported that serum and urine L-FABP may be a new diagnostic marker for liver damage in patients with both acute and chronic hepatitis C infection (Cakir et al., 2017). Interestingly, in Atlantic salmon challenged with *P. salmonis*, L-FABP was up-regulated in resistant families and simultaneously down-regulated in susceptible families (Pulgar et al., 2015), suggesting a transcriptional regulation in response to *P. salmonis* infection and a putative expression marker of resistance to SRS.

Genes coding NACHT, LRR and PYD domains-containing protein 12 (NLRP12); NACHT, LRR and CARD domains-containing protein 3 (NLRC3); voltage-dependent L-type calcium channel subunit beta-2 (CACNB2); heparan sulfate glucosamine 3-O-sulfotransferase 3A1 (HS3ST3A1) and histidine triad nucleotide-binding protein 1 (HINT1) were also selected as candidate genes for SRS resistance. NLRP12 and NLRC3 are two cytosolic proteins that share two functional domains (NACHT and LRR). NLRP12 was one of the best ranked genes, adjacent to the leader SNP and adjacent to more than one SNP simultaneously for DD in AS. This protein functions as an attenuating factor of inflammation in monocytes, by negative regulation of the NF-κB activation (Fata et al., 2013). In murine macrophages, a significant expression increase has been shown in cells infected with the intracellular parasite *Leishmania major* compared to non-infected macrophages (Fata et al., 2013). NLRC3 is also a negative regulator of the innate immune response mediated by the inhibition of Toll-like receptor (TLR)-dependent activation of the transcription factor NF-κB (Schneider et al., 2012). The presence of these genes suggests that the control of the inflammatory reaction in response to *P. salmonis* infection could be essential to combat SRS.

Finally, some members of heparan sulfate glucosamine 3-O-sulfotransferase (like HS3ST3A1) have been shown to mediate the herpes simplex virus type-1 (HSV-1) entry and spread in zebrafish (Antoine et al., 2014). Also, some members of CACNB2 changed their expression levels in response to *P. salmonis* in multiples tissues of Atlantic salmon (Tacchi et al., 2011), while HINT1 responds transcriptionally to *Salmonella typhimurium* and infectious pancreatic necrosis virus (IPNV) infection in zebrafish and Atlantic salmon, respectively (Stockhammer et al., 2010; Robledo et al., 2016).

To the best of our knowledge, this is the first time that functional candidate genes underpinning resistance to *P. salmonis* are proposed based on a comparative genomics approach comparing GWAS results for the same trait in different fish genus/species. We hypothesize that variations in the sequences of these genes could play important roles in the host response to *P. salmonis* infection, which could be tested through new genetic approaches such as gene editing using CRISPR-Cas9, and utilized through genomic selection or more traditional selection practices. All this information together can be used to generate better control and treatment measures for one of the most important bacterial disease affecting salmon aquaculture.

## Conclusions

Although *P. salmonis* resistance has previously been described as polygenic trait, our comparative genomics approach based on GWAS results for the same trait in different salmonid species allowed us to identify around one hundred candidate genes that may explain resistance to *P. salmonis*. Of these, 21 are suggested to be strong functional candidates influencing the trait. These genes are associated with multiple biological processes, including: dependent on kinase activity, GTP hydrolysis, helicase activity, lipid metabolism, cytoskeletal dynamics, inflammation and the innate immune response. We hypothesize that variations in the sequences of these genes could play an important role in the expression and/or activity of their encoded proteins and consequently in the resistance to *P. salmonis*. This information could be used to generate better control and treatment measures, based on selective breeding or new drug development, for one of the most important bacterial disease affecting salmon aquaculture.

## Data availability

Genotype and phenotype data for Atlantic salmon and rainbow trout are available on the online digital repository figshare figshare.com/s/e70a3a84c05ea60e1074 and figshare.com/s/221a39319b236d46f9fc respectively. Coho salmon genotype and phenotype are at available doi.org/10.5061/dryad.b273q6p). and at 10.6084/m9.figshare.7886183.

## Acknowledgements

Aguas Claras, Pesquera Antares and Salmones Chaicas provided the CS, RT and AT datasets, respectively. Thanks to FAPESP (2014/20626-4; 2015/25232-7) and CNPq (308636/2014-7) for financial support. AB and KC acknowledge the National Commission of Scientific and Technologic Research (CONICYT) for the funding through the National PhD funding program. RP acknowledge the National Commission of Scientific and Technologic Research (CONICYT) for the funding through the Fondecyt program (11161083). AB acknowledges the Government of Canada for the funding through the Canada-Chile Leadership Exchange Scholarship. We also want to thank the World Congress on Genetics Applied to Livestock Production (WCGALP) as this work has been partially presented on this conference (Yoshida et al. 2018b)

## Ethics approval and consent to participate

All experimental challenges and sampling procedures were approved by the Comité de Bioética Animal from the Facultad de Ciencias Veterinarias y Pecuarias, Universidad de Chile (Certificate N08-2015).

## Authors’ contributions

JMY conceived of and designed the study and drafted the manuscript. GrY assessed the GWAS analyses. AP and RP assessed the comparative genomic analyses and contributed with the first draft of the manuscript and discussion. LB, MEL and KC contributed with the RT and AS sampling, genotyping and quality controls. AB performed DNA extraction from CS samples, contributed with initial draft of the manuscript and performed library construction. KrC performed ddRAD library construction and assessed the comparative sequences analyses between species. RC and JPL contributed with study design, analyses and discussion. All authors have reviewed and approved the manuscript.

---

**Figure S1.**
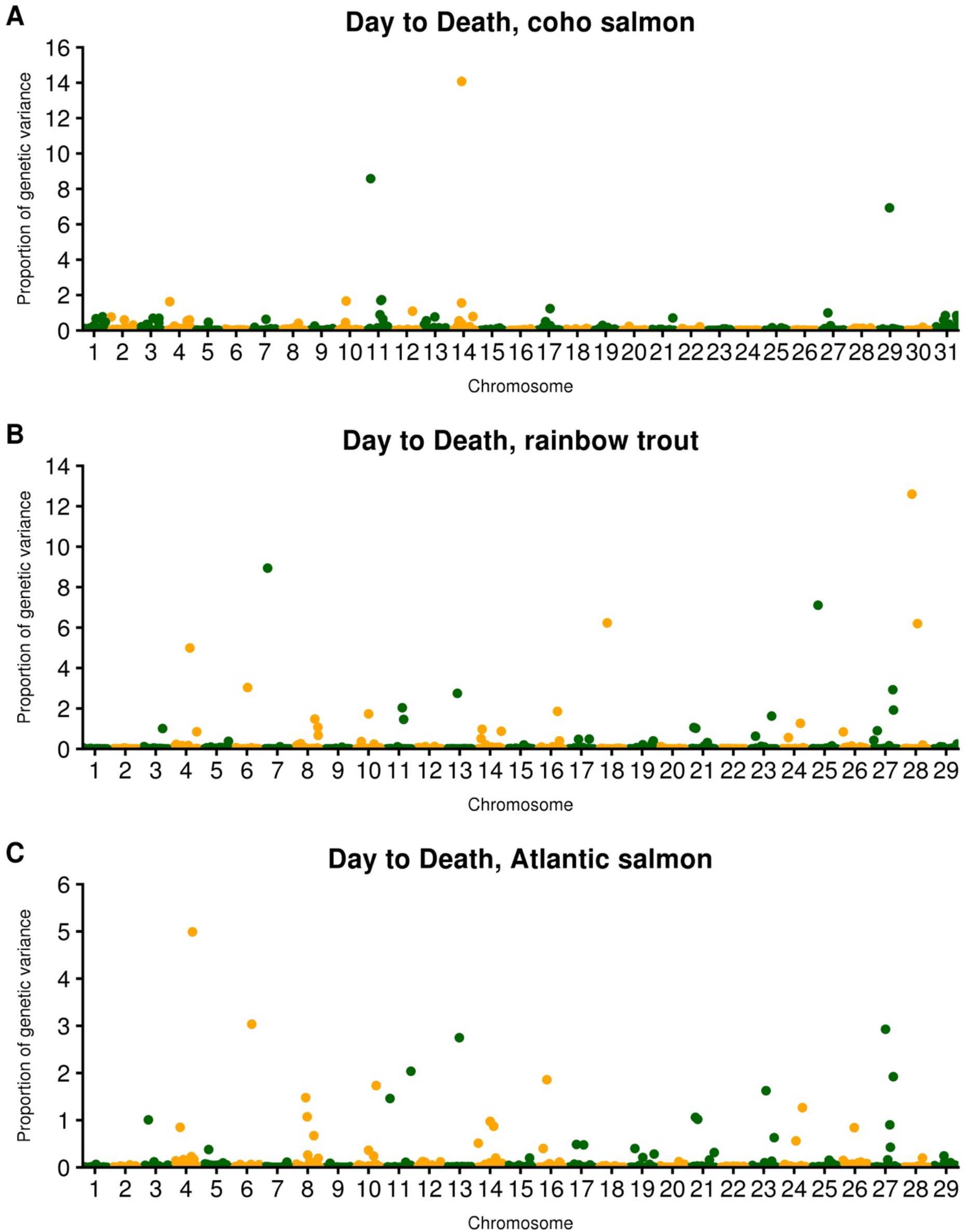
Manhattan plots for resistance to *P. salmonis* measured as day of death (DD) in coho salmon (CS), rainbow trout (RT) and Atlantic salmon (AS). Y-axis represents percentage of the genetic variance explained by each marker.

**Figure S2.**
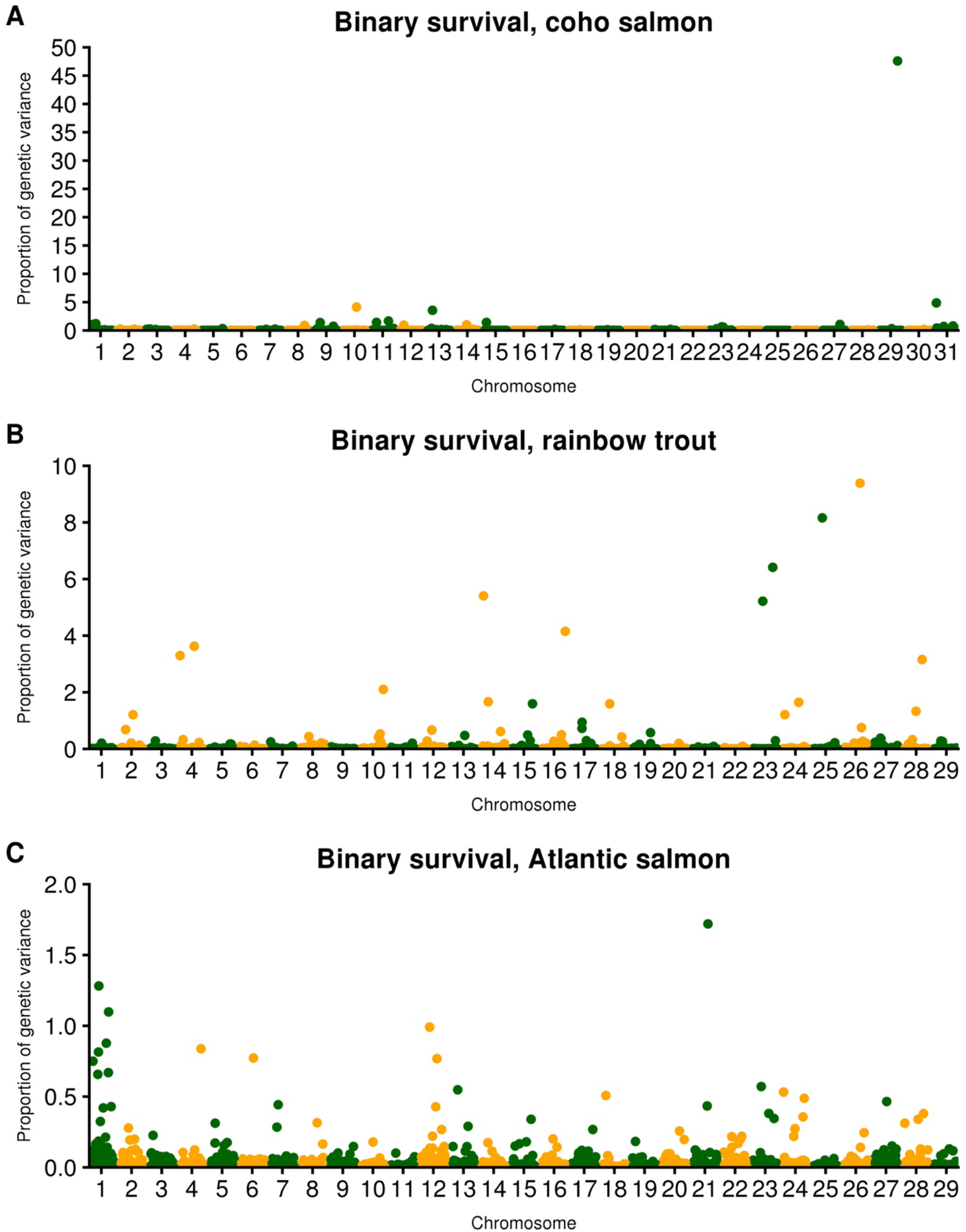
Manhattan plots for resistance to *P. salmonis* measured as binary survival (BS) in coho salmon (CS), rainbow trout (RT) and Atlantic salmon (AS). Y-axis represents percentage of the genetic variance explained by each marker.

**Figure S3.**
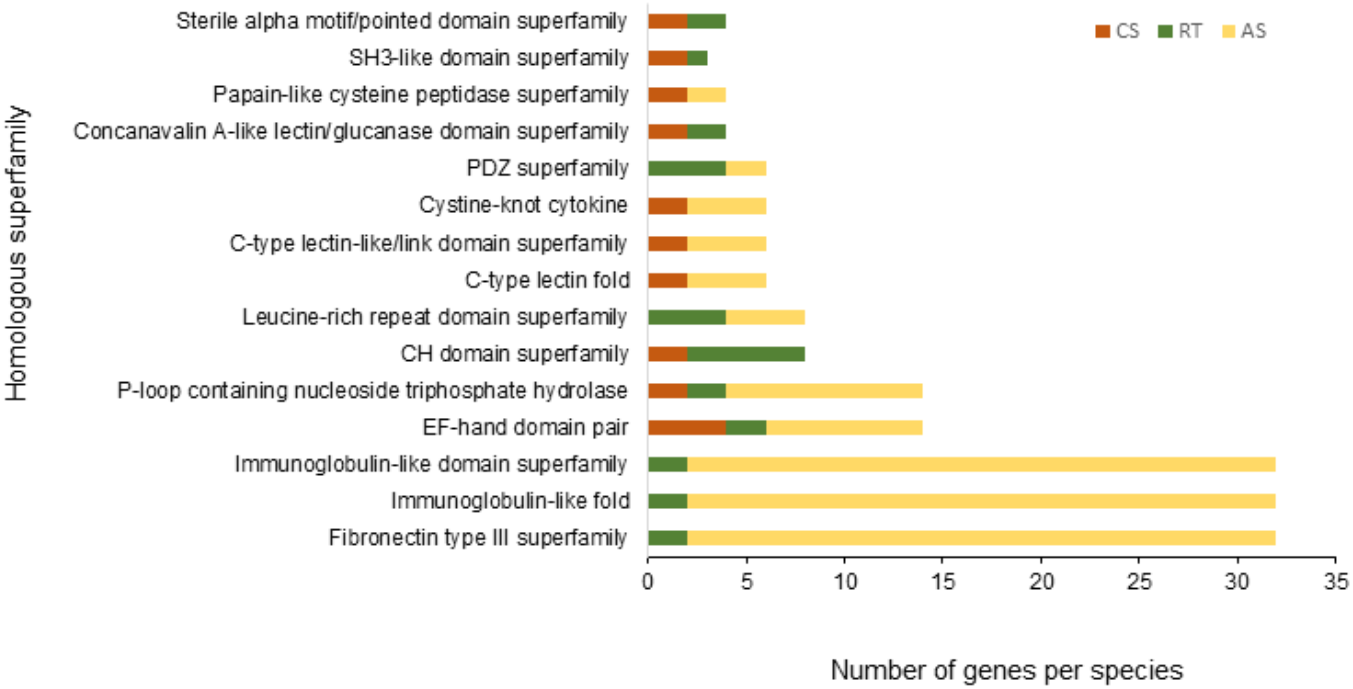
Homologous superfamilies (InterPro) associated to 100 random selected proteins from coho salmon (CS), rainbow trout (RT) and Atlantic salmon (AS) genomes.

